# Using UAV-Imagery for Leaf Segmentation in Diseased Plants via Mask-Based Data Augmentation and Extension of Leaf-based Phenotyping Parameters

**DOI:** 10.1101/2022.12.19.520984

**Authors:** Abel Barreto, Lasse Reifenrath, Richard Vogg, Fabian Sinz, Anne-Katrin Mahlein

## Abstract

In crop production plant diseases cause significant yield losses. Therefore, the detection and scoring of disease occurrence is of high importance. The quantification of plant diseases requires the identification of leaves as individual scoring units. Diseased leaves are very dynamic and complex biological object which constantly change in form and color after interaction with plant pathogens. To address the task of identifying and segmenting individual leaves in agricultural fields, this work uses unmanned aerial vehicle (UAV), multispectral imagery of sugar beet fields and deep instance segmentation networks (Mask R-CNN). Based on standard and copy-paste image augmentation techniques, we tested and compare five strategies for achieving robustness of the network while keeping the number of labeled images within reasonable bounds. Additionally, we quantified the influence of environmental conditions on the network performance. Metrics of performance show that multispectral UAV images recorded under sunny conditions lead to a drop of up to 7% of average precision (AP) in comparison with images under cloudy, diffuse illumination conditions. The lowest performance in leaf detection was found on images with severe disease damage and sunny weather conditions. Subsequently, we used Mask R-CNN models in an image-processing pipeline for the calculation of leaf-based parameters such as leaf area, leaf slope, disease incidence, disease severity, number of clusters, and mean cluster area. To describe epidemiological development, we applied this pipeline in time-series in an experimental trial with five varieties and two fungicide strategies. Disease severity of the model with the highest AP results shows the highest correlation with the same parameter assessed by experts. Time-series development of disease severity and disease incidence demonstrates the advantages of multispectral UAV-imagery for contrasting varieties for resistance, and the limits for disease control measurements. With this work we highlight key components to consider for automatic leaf segmentation of diseased plants using UAV imagery, such as illumination and disease condition. Moreover, we offer a tool for delivering leaf-based parameters relevant to optimize crop production thought automated disease quantification imaging tools.

## 1 Introduction

Imaging sensors attached to unmanned aerial vehicles (UAVs) are systems that currently revolutionize the way to monitor agricultural fields [1]. Agricultural management practices are feasible after analysis of high spatial and temporal resolution images, allowing decision making at the right place, with the right intensity, and at the right time. The list of practical applications of UAV-systems starts with the area of plant breeding and phenotyping, and has further applications for plant protection, yield prediction, growth vigor, nutrient status, weed detection and drought stress. Moreover, UAV-systems support precision agriculture because crop production is optimized by maintaining or increasing yield, while reducing environmental impact and resources for pest, water and nutrient management [2].

Disease quantification is a common visual scoring activity for plant breeding and decision making in plant protection nowadays. This activity must be executed several times over vegetation development by appropriately trained staff or experts [3]. The most established parameters for quantification are disease incidence (*DI*) and disease severity (*DS*) [4, 5]. For phenotyping *DI* and *DS* in sugar beets, experts collect a representative number of sugar beet leaves, which are assessed for presence or absence of symptoms, and for determining the degree of damage [6]. According to Reynolds et al. [7], phenotyping work like visual scoring represents the major proportion of costs in experimental trials. Variability and repeatability of collected data is also an issue in field assessments: “Inter-rater” source of error is introduced when different individual experts are required for phenotyping key disease development stages. This error source caused by the high manpower need is a disadvantage for extensive experimental fields. Similarly, other factors negatively influence the variability and repeatability of observations, including the effect of noise, heat, exhaustion or time allocated for an assessment [5]. All those factors emphasize the limitations of visual scoring methods and motivate the development of innovative and automated UAV-based imaging approaches for quantification of plant diseases [8, 9, 10, 11].

In past studies used RGB and multispectral UAV-systems for disease quantification in sugar beet fields. Jay et al. [8] and Görlich et al. [9] segment limits of plot regions from RGB and multispectral orthomosaic images to calculate *DS* in variety trials. Similarly, a pipeline by Günder et al. [12] to segment individual plants, was extended for an application in disease quantification of Cercospora leaf spot (CLS) to classify infested plants according their disease categories [10]. Circular regions within a plot are considered for an automated analysis of multispectral images to calculate *DI, DS*, and additional parameters such as area of foliage, area of healthy foliage, number of lesions and mean area of lesion by unit of foliage [11]. Whether circular-, plant-, or plot-based regions, this image-based scoring shows robustness to calculate *DS*. Nevertheless, *DI* is an aspect to improve: Barreto et al. [11] highlight the disadvantages of delimiting scoring in circle regions within a plot. False positive pixel classification of non-diseased regions leads to inaccurate quantification of diseased units for *DI*. Visualization of scored multispectral images shows that the principal reason of disease misclassification is the pixel quantification of harvest residues of previous crop laying on soil regions. This highlights a potential application of leaf segmentation because image-regions with high misclassification rate are removed from determining *DI*. While leaf segmentation could also contribute to *DS*, this has not been tested yet.

Image-based leaf phenotyping requires detecting and delineating a representative number of leaves for a later parameter calculation. Sugar beet canopy is a complex structure of non-uniform leaves. Individual leaves are positioned with extreme overlap, mutual occlusion, at different heights and diverse orientation. Moreover, leaf appearance is dynamic, either by natural develop stages and senescence, or by exogenous factors like diseases. Leaves change in size, and color from green to yellow, by degradation of chlorophyll content, and later from yellow to brown when necrotic tissue dominates the canopy. At last, weather conditions play also role. UAV monitoring activities must be able to cope with cloudy and sunny sky conditions if farmers are to obtain an on-time decision. Furthermore, shaded regions caused by passive illumination have to be considered for image-based individual leaf segmentation.

To solve the task of leaf identification and segmentation from UAV-data, adequate data analysis approaches are required. This task can potentially be automated by deep learning models in computer vision, more specifically instance segmentation [13]. Unfortunately, the main limitation of deep learning models in agriculture is the need for a high number of labeled images [14]. In the context of UAV-data of agricultural fields, labeling individual leaves in real images is time intensive, making this work the bottleneck for the availability of labeled data.

In this paper, we make two main contributions: (i) we address the challenge of leaf segmentation from multispectral UAV-based images with a limited number of labeled images. We evaluate augmentation approaches including basic image manipulation and copy-paste techniques to create a data set of adequate size for training instance segmentation models (Mask R-CNN). For this specific aim, we consider recommendations as described by Kurnichov et al. [15] using a copy-paste data augmentation approach for banana plantain and arabidopsis images. We identify the best model by testing performance under diverse disease and illumination conditions. (ii) We apply our best leaf segmentation model to large orthomosaic images, and integrate Mask R-CNN into a pipeline extend the number of parameters for disease quantification beyond the circle-based parameters. Lastly, we apply the pipeline in a variety trial and evaluate the performance by comparing expert with automated scored data of *DS* collected in time-series.

## 2 Field monitoring and methodology

The monitoring campaign took place from 2019 to 2021 in four locations [16, 11]. We use multispectral UAV-systems: a DJI Inspire 2 with a 5-channel multispectral sensor, Micasense RedEdge-M; and a DJI Matrice 210 with a the 6-channel sensor variant, Micasense Altum. To get high resolution images, flight missions were planned to deliver a ground sample distance (GSD) between 2.5 and 4.1 mm. We truncate and calibrate the raw images from digital numbers to reflectance according to Barreto et al. [11]. For the photogrammetry, we use the software Agisoft Metashape Professional for stitching of truncated multichannel images, and export the output as orthomosaic. We generate image patches to cover a field area of 1.28 × 1.28 m and scale these to 512 × 512 px by resampling (Fig. 1a). We use five and six channel re-scaled arrays for defining RGB composite images from reflectance values by using *V* = max(*λ*_*BLUE*_, *λ*_*GREEN*_, *λ*_*RED*_), and the following formulae:

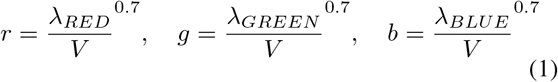

**Figure 1:**
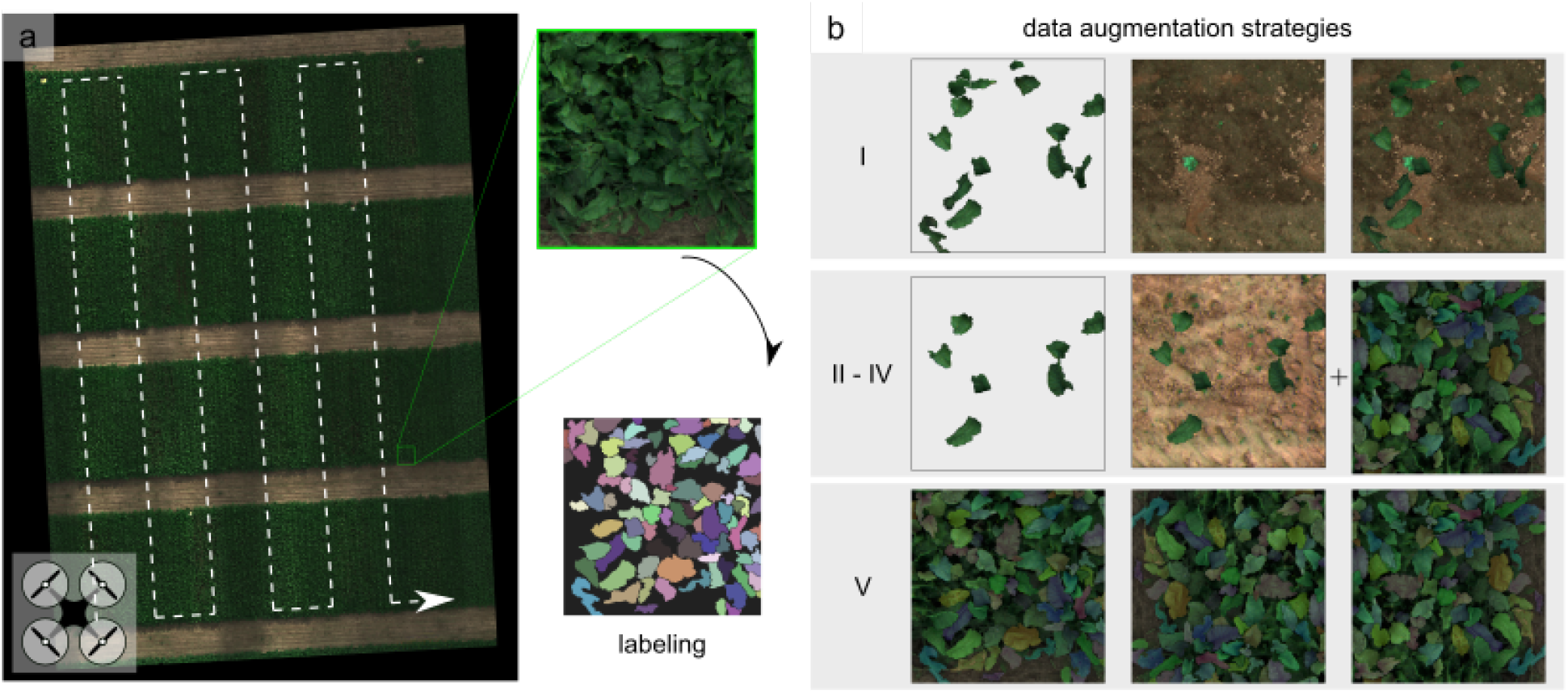
a) Experimental field trial with flight mission, RGB composite orthomosaic, and labeling of image patches (left). b) Data augmentation following five strategies an (I) exclusively copy-paste technique, (II-IV) combination of copy-paste and simple techniques, and (IV) exclusively simple techniques.

### 2.1 Modeling leaf segmentation

#### 2.1.1 Data augmentation

Manual data labeling is a very difficult and time-consuming task, particularly for sugar beet images which have highly overlapping leaves with almost identical colors. We compare five strategies (see Fig. 1b) to generate sufficient training data while keeping the labeling efforts within reasonable bounds. First, we completely label every leaf in 100 images (512 × 512 pixels) containing healthy and diseased sugar beet plants under sunny and cloudy light conditions. In total, the images contain 2951 leaves without occlusion and we split the individual leaves 80/20 into training and validation set. Eighteen additional completely labeled images serve as a hold-out test set to evaluate the performance of different models. For the background, we select 758 background UAV-images (without sugar beet plants), containing different field conditions such as: diverse soil humidity, weed pressure and soil tillage.

We use the training data and background images in four of five data augmentation strategies (I-V). For all strategies the data sets contain 10000 images (training to validation ratio of 8000 to 2000). For strategy I we use a copy-paste augmentation technique. For this, we randomly position 70-140 segmented leaves onto randomly selected background images. We use color jittering, rotation, and flipping on each leave image before pasting it to the background. Splitting the leaves beforehand into training and validation set avoids data leakage. strategies II to IV are mixed strategies. We create a proportion of the images with the copy-paste method described above. Additionally, we add the 100 completely labeled images, and use standard image augmentation methods: flip, rotation, channel shift, crop and brightness shift. For strategy II, we use the 100 original images and create 4900 copy-paste images. Then, we augmented the whole set two times with standard augmentations to obtain 10000 images. Strategy III uses 100 original images and 1900 copy-paste images, and we augment with basic augmentations five times. For strategy IV, we use 100 original images and 100 copy-paste images and fifty times basic augmentations. The last strategy (V) uses only basic manipulation, where each original image was augmented 100 times.

#### 2.1.2 Training

For instance segmentation, we use a Mask R-CNN architecture [17] with a ResNet-101 backbone for ROI (region of interest) prediction. We train the models on four RTX 5000 GPUs running in parallel and use the adapted Mask R-CNN implementation for Tensorflow 2.0 [17] with modules Cudnn 7.6.5.32-10.2, cuda 10.1.243 and Tensorflow 2.2.0. Before training, we specify the classes “leaf” and “background”. We put the number of ROIs to train per image from 200 up to 256 and anchors to train per image from 256 up 512. We initialize our models with weights pretrained on the COCO data set [13]. Training the model for one step takes about 22 seconds. Since we make 500 steps per epoch we get a training time of three hours for one epoch. Mask R-CNN implements a multi-task loss, ℒ= ℒ_*cls*_ + ℒ_*box*_ + ℒ_*mask*_, to balance correct class predictions, accurate bounding boxes and exact masks. In this study, we train all models for 50 epochs. During the first 25 epochs, the learning rate is kept at 0.001, then changed to 0.0001 until the end of the training.

#### 2.1.3 Evaluation and metrics

We group test set images in six categories based on illumination conditions (cloudy and sunny), as well as the degree of disease damage (healthy, medium and severe). We evaluate the performance of the models with precision, recall and average precision (AP). Precision measures how many of our predicted leaves are actually leaves, while recall evaluates how many of the actual true leaves are detected by the model. Precision and recall depend on a threshold measuring intersection over union (IoU) between a predicted and a ground truth leave. We set this threshold to 50% for precision and recall. This means that to accept a prediction as correct it needs to overlap at least to 50% with a ground truth leave. The second hyperparameter is the confidence score threshold describing how confident the model must be to use the leave prediction. In this paper, we compare values of 0.5 and 0.75 for the confidence score (CS).

For AP, we measure precision and recall for all possible confidence thresholds and plot them in a curve. The area under this curve is the AP. It is large when the true predictions come with high confidence values and there are few false positives.

#### 2.1.4 Leaf segmentation results

In this section we compare the five augmentation strategies for leaf segmentation in diseased sugar beet plants. In Table 1 the AP, precision and recall under two confidence scores CSs are compared. In all approaches, by increasing the CS value, the number of false negative or missed leaves will increase. In contrast to recall, precision increases with higher CS. Increasing CS from 0.50 to 0.75 yields between 6% to 12% better precision results. The advantages of models with CS of 0.50 was reflected on a higher AP in comparison to models with 0.75 CS. Overall, the best AP was found in the model of strategy V with a value of 0.31. This strategy consist of augmenting fully labeled images only using standard techniques to increase the train and validation set to 10000 images. In contrast to the study by Kuznichov et al. [15] on avocado plants, we do not find that a copy-paste augmentation strategy improves our results.

**Table 1:** Error metrics of Mask R-CNN models for augmentation strategies I to V considering two confidence scores for prediction in the test set

|  | Strategy | I | II | III | IV | V |
| --- | --- | --- | --- | --- | --- | --- |
| CS | 0.50 |  |  |  |  |  |
|  | AP | 0.036 | 0.161 | 0.251 | 0.297 | **0.305** |
|  | Precision | 0.377 | 0.457 | 0.568 | 0.623 | 0.642 |
|  | Recall | 0.064 | 0.249 | 0.336 | 0.384 | 0.402 |
|  | 0.75 |  |  |  |  |  |
|  | AP | 0.015 | 0.102 | 0.162 | 0.183 | 0.162 |
|  | Precision | 0.436 | 0.581 | 0.695 | 0.726 | 0.723 |
|  | Recall | 0.022 | 0.139 | 0.201 | 0.221 | 0.200 |

CS= confidence score.

Image context is also an aspect to consider in the leaf segmentation problematic. This plays a role in object detection and segmentation, when the object size is small in relation to complete image [18]. Image context can be ordered in three perspectives: local context, global context, and context interactives. The epidemiological development of the leaf disease and the nadir UAV-perspective by imaging should be an example of image context for the used data augmentation techniques, because specific enviromental conditions and coexist with other related objects are not emulated. A previous study was able to achieve better performance when image context is simplifyed. If sugar beet plants are recorded before canopy closure and without disease pressure, when less number of leaves and overlapping as well as no leaf senescence factor is visual; the AP value can be improved to 0.413 for using Mask R-CNN [19].

#### 2.1.5 Environmental conditions for monitoring diseased plants

We test on data sets with categories “cloudy” and “sunny” separately to further explain the performance of a Mask R-CNN leaf segmentation model under two different environmental illumination conditions. For this evaluation, the best model in terms of AP (strategy V) was analyzed in Table 2 for all six categories. Data recorded under sunny illumination conditions is the principal source for decrease of performance. UAV-images of sunny days drop in AP of 7% against cloudy and diffuse illumination. Nevertheless, it remains still the question if drop in performance is mainly affected by false positive or false negative detections. Analyzing in deep a further parameter for metric such as precision, it is possible to observe that this parameters has a more intensive drop in performance in comparison with AP while recall keeps on the same levels (drop in precision of 12.3%). We conclude that analyzing images recorded under sunny illumination conditions increases the number of false positives or objects incorrectly detected as leaves. Illumination is an image condition that affects intra-class variation. This variation drastically impacts the performance of object detection deep learning approaches, because object appearance change in brightness and shading [20]. Furthermore, the effect of degree of disease damage is not clearly visible (Table 2). This last phenomena needs to be considered in context a practical application for leaf instance segmentation.

**Table 2:** Performance of model from strategy V in test set considering patch-based image annotation for two light conditions and three disease severity stages

|  |  | Healthy | Middle | Severe | Average |
| --- | --- | --- | --- | --- | --- |
| Cloudy | AP | 0.315 | 0.350 | 0.354 | **0.339** |
|  | Precision | 0.741 | 0.670 | 0.701 | 0.704 |
|  | Recall | 0.380 | 0.441 | 0.436 | 0.419 |
| Sunny | AP | 0.304 | 0.265 | 0.242 | **0.270** |
|  | Precision | 0.680 | 0.535 | 0.528 | 0.581 |
|  | Recall | 0.399 | 0.404 | 0.350 | 0.384 |
| Average | AP | **0.309** | **0.307** | **0.298** | 0.305 |
|  | Precision | 0.710 | 0.602 | 0.615 | 0.642 |
|  | Recall | 0.390 | 0.423 | 0.393 | 0.402 |

### 2.2 Application of leaf segmentation for UAV-based parameter extraction

#### 2.2.1 Disease quantification in variety trial

We explore the potential of leaf segmentation for monitoring field experiments in a variety trial in 2021 near Göttingen, Germany. The goal of this experiment was to quantify resistance against the disease Cercospora leaf spot (CLS). We arranged a plot-trial with five sugar beet varieties and two fungicide strategies in a two-factorial block design with four blocks or repetitions. The two fungicide strategies are: control with fungicide, and inoculated without fungicide. The registration identifiers of the five sugar beet varieties *Beta vulgaris* L. ssp. *vulgaris* (A-E) are: 3012, 2444, 3290, 3316 and 3706 respectively. Selected varieties belong to the national variety list of the German Federal Plant Variety Office (Bundessortenamt) [21].

We carry out visual scoring of *DS* of CLS as ground truth simultaneously to the UAV flights, and quantify symptoms as an average value of the plot by assessing middle leaves in percentage estimating a representative infected from total leaf area [6]. The assessment is conducted at leaf level. In total, we randomly sample 100 leaves (25 leaves/plot) per treatment and inspect them for CLS symptoms.

#### 2.2.3 Prediction on large images

Orthomosaic images from UAV-imagery outrun current GPUs capacity for prediction. To extend the model application to a large image from a complete mapped field, we use a sliding windows procedure (Fig. 2). For this purpose and as proposed by Machefer et al. [22], we crop the RGB composite orthomosaic image (*I*) in patches with an overlap (15 cm or 60 px, *o*) between neighbours. We use the segmentation model to predict leave instances sequentially on each patch; and eliminate instances with 100% cover in the overlap region. After selection, we reconstruct the remaining leaves in a new orthomosaic image (*I*_*leaf*_). Algorithm 1 shows the procedure.

**Figure 2:**
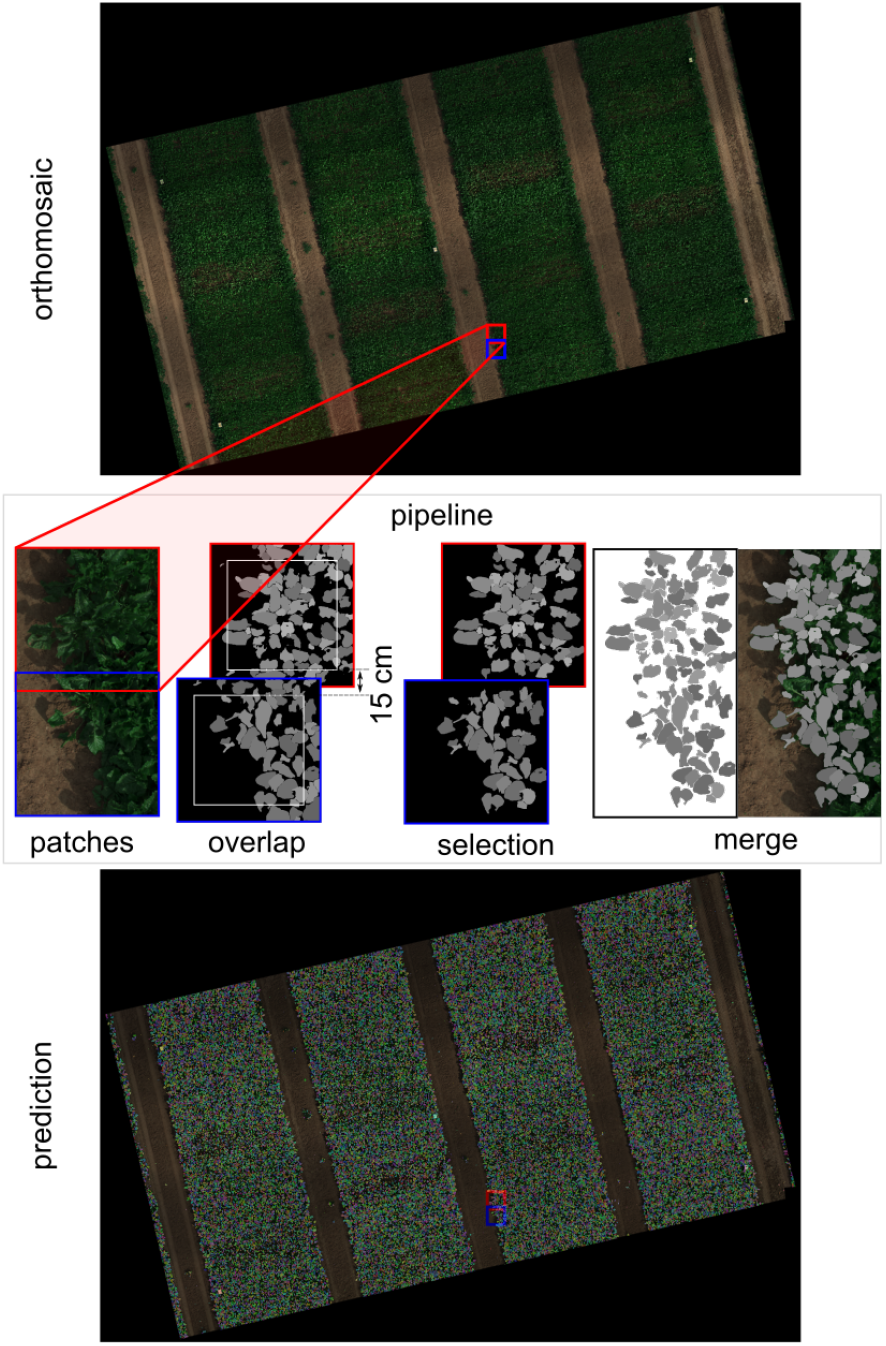
Pipeline sliding windows for Mask R-CNN prediction of large orthomosaic images, considering an overlap between windows of 15 cm.

##### Algorithm 1

Implementation of sliding window prediction for large images

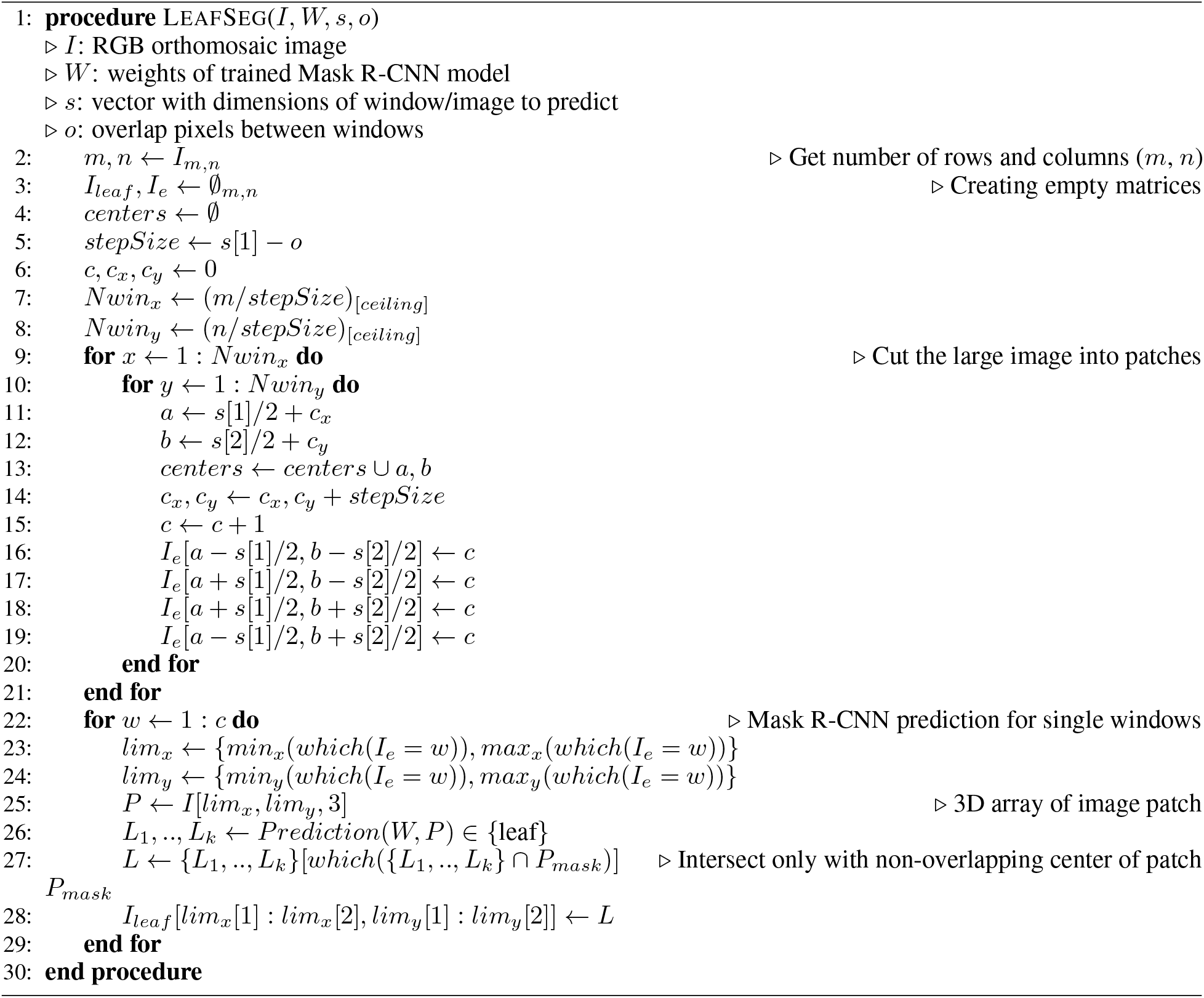

#### 2.2.3 Instance segmentation for leaf parameter extraction

The availability of *I*_*leaf*_ increases the potential for extraction of plant phenotyping parameters. In this study, we propose the integration of this output to a pipeline for pixel-wise classification and extraction of disease-relevant parameters as specified by Barreto et al. [11]. The new pipeline starts with radiometric calibration and photogrammetric processing of raw images resulting in the multispectral orthomosaic and the digital surface model (DSM). Then, we create a RGB composite orthomosaic image as specified in Section 2 (Figure 3a). In the next step, prediction of large image takes place to deliver the *I*_*leaf*_ output (Figure 3b). We calculate image features of multispectral orthomosaic and DSM are store them for later feeding them to two multiclass classifiers, a partial least squares discriminant analysis (PLS-DA), and a support vector machine radial (SVMR). At this level each pixel (*Z*) is assigned one of four classes, “other”, “soil”, “diseased”, “healthy”, *Z* ∈ {*other, soil, diseased, healthy*}. This results in a binary array of four layers or images: *O*_*i,j*_, *S*_*i,j*_, *D*_*i,j*_, and *H*_*i,j*_, assigned to the four classes respectively. The overlay of any of those outputs with an instance *L* (*L* ∈ *I*_*leaf*_) returns four outputs with the delimited instance region: *O*_*L*_, *S*_*L*_, *D*_*L*_, and *H*_*L*_ (Figure 3c). The binary image *D*_*L*_ is relevant for disease quantification; therefore, clusters (*c*) can be extracted from *D*_*L*_ (Fig. 3c) by labeling eight connected cluster pixels [23].

**Figure 3:**
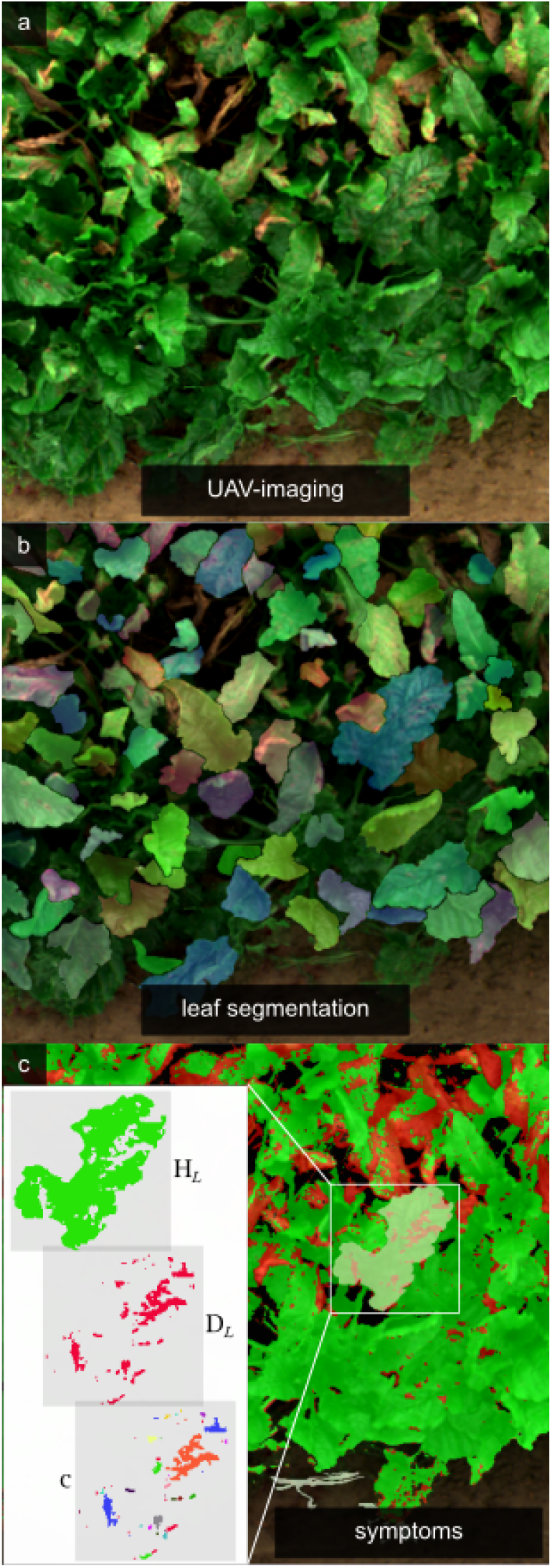
Prediction of disease quantification parameters by application of leaf segmentation. a) RGB composite of diseased plants, b) segmentation of leaves, and c) multiclass pixel-classification and parameter extraction of leaf instances as healthy region (*H*_*L*_), diseased region (*D*_*L*_), and number of clusters within a leaf (*c*).

Considering the image area (*A*_*L*_) and image slope (ζ_*L*_) as calculated features from DSM within an instance *L* [24, 11], we calculate the leaf-based parameters with the formulae in table 3. These parameters are: leaf area (*A*_*l*_), leaf slope (ζ_*l*_), diseased leaf area (*A*_*D*_), healthy leaf area (*A*_*H*_), disease severity (cover based, *ds*_*l*_), disease severity (area based, *DS*_*l*_), number of clusters (*c*), and mean cluster area 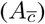.

**Table 3:**
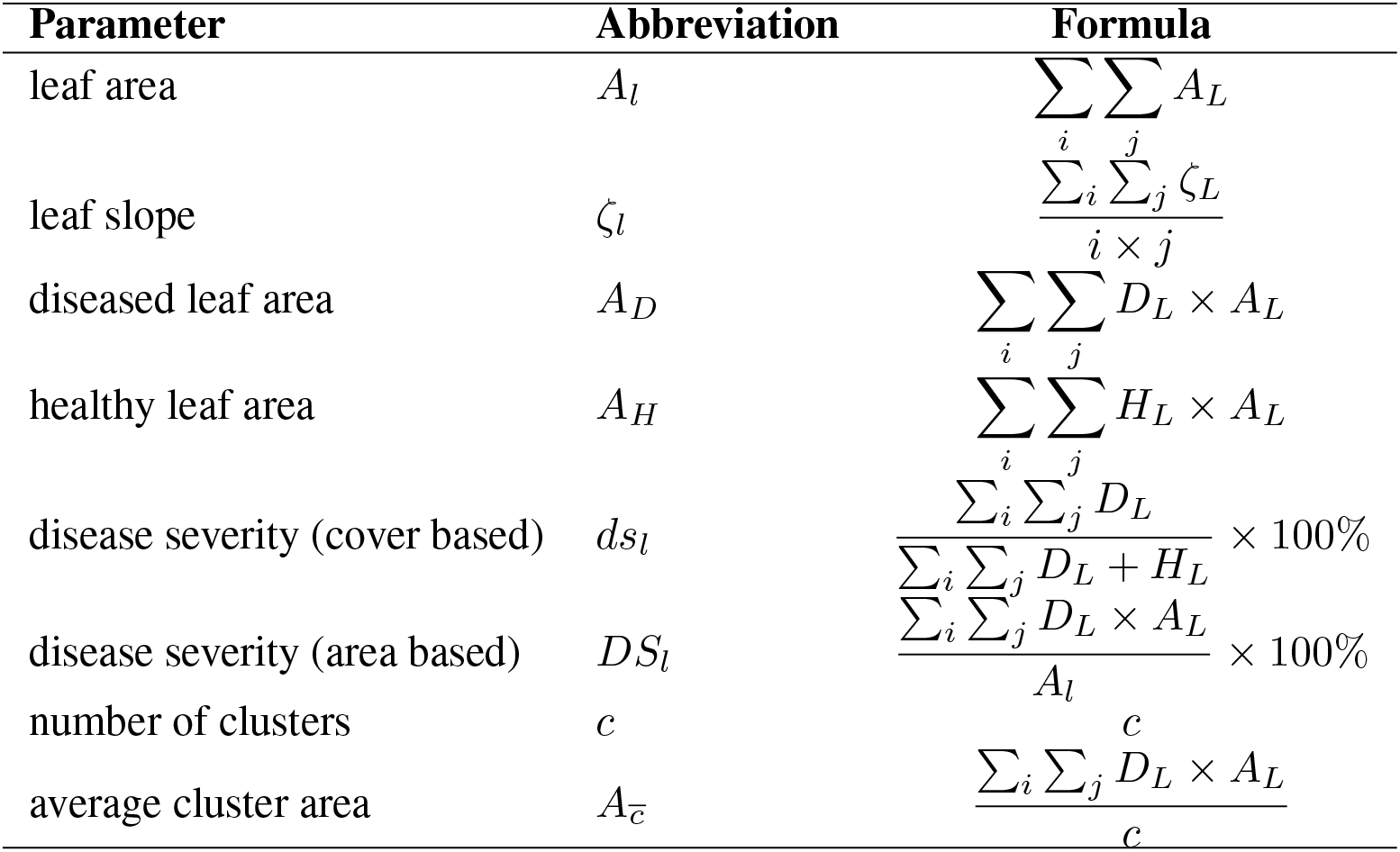
Definition for parameters inside leaf instance

#### 2.2.4 Automatic scoring at plot level

Some parameters can not be directly determined from one instance, but they require a sample of instances within a field area. In field trials, this sampling takes place within a plot. A representative sampling for monitoring diseases normally consists of taking 100 leaves for scoring per plot [6]. In the same way, we can express the parameters mentioned in table 3 as an average within the field area. Moreover, we can calculate the relevant parameter *DI* in a plot-based manner. To determine *DI*, it is necessary to define an affected or diseased unit as mentioned by [25, 26, 11]. In the past, we defined an instance as diseased if at least one lesion was present or if the pixel summation in *D* was greater or equal than one cluster (*c* ≥ 1). We can also define a diseased unit with another parameter such as *ds*_*l*_. Similarly, it is possible to establish threshold values to order diseased leaves (Fig. 6a).

#### 2.2.5 Leaf segmentation for disease severity

We compare the disease quantification parameter, *DS* from UAV-based and expert-based sources in figure 4. Figure 4a and b show a parabolic behavior of the UAV-based *DS* for models with the lowest and the highest AP (strategy I and V). This behaviour presents a maximum value of UAV-based *DS* at 40% of the ground truth data, and seems to be a limit for the automatic calculation of leaf-based *DS*. By evaluating the proportion of variance in linear regression (*r*^2^) between both *DS*s, we observe slightly better results in the linear regression model of the UAV-based data from strategy V compared to strategy I. Moreover, there are no clear differences between models with CS of 0.50 or 0.75.

**Figure 4:**
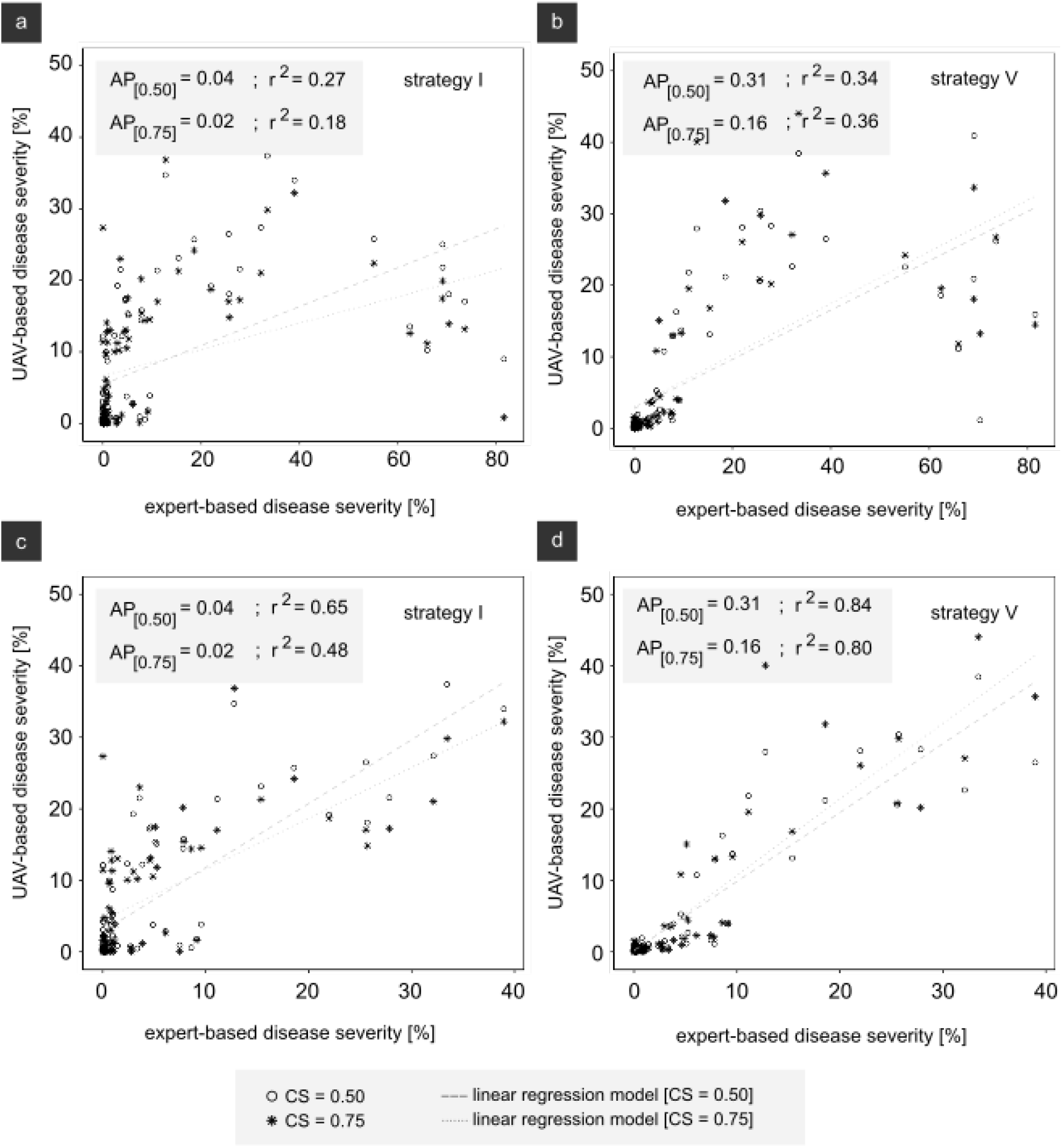
Relationship of unmanned aerial vehicle (UAV)- and expert-based scores for confidence scores (CS) of 0.50 and 0.75 in two monitoring dates of experimental test field. Comparision of all ground-truth values for model in data augmentation a) strategy I and b) strategy V. Comparison for ground truths lower than 40 % in c) strategy I and d) strategy V. Monitoring dates: 17th and 31st August 2021.

We attribute this parabolic relationship of UAV-based *DS* to the image context and the combined effect from nadir UAV perspective and the new leaf growth emerging of a typical severe epidemics of CLS [27]. The nadir perspective of UAV record mainly above-located leaves which are under severe epidemics and covered by healthy new leaves. This entails special advantage of visual scoring because from a side perspective heavy diseased and small leaves are quantified with high *DS* values [28], while low *DS* values are delivered from UAV-based scores by quantifying healthy new leaves.

To mitigate the effect of image context by analyzing less disease development stages, in Fig. 4c and d were deleted all data with high expert-based values (*DS* higher than 40%). In general, values of *r*^2^ increase significantly after data deletion. Here, the model with the highest AP (strategy V) shows a clear advantage against model of strategy I. In addition, the model with CS of 0.75 performs slightly better than the model with CS of 0.50. A possible solution against this image context challenge is to introduce a new class of leaf instances for leaves with damage higher than 40% in order to give priority to the detection of this morphological different leaves.

#### 2.2.6 Epidemiological development of disease quantification parameters

In this section we show one of the principal applications of leaf segmentation for parameter extraction and plant phenotyping of variety trials for quantitative resistance quantification. As described in Section 2.2.1, we work with the experimental design with five varieties and two fungicide strategies. In figure 5, we show the development of each genotype during the complete period for disease monitoring with average values of *DS*_*l*_, *A*_*l*_, and *c*. We compare the development of *DS*_*l*_ in figure 5a and b for the control with fungicide and inoculated without fungicide variants. Here we see the effect of disease pressure by applications of fungicide, where the control with fungicide variant keeps all varieties healthy until the beginning of September (figure 5a). We further observe resistant characteristics of a genotype with high disease pressure. In figure 5b, variety A shows the most susceptible characteristics, and variety E presents apparently the highest tolerance against CLS. The leaf area is also affected by the genotype and disease pressure (Fig. 5c and d). In Fig. 5c we see that variety C presents the smallest *A*_*l*_ until mid August, while varieties D and E show the biggest *A*_*l*_ during the complete monitoring period. *A*_*l*_ is further affected by disease pressure. UAV-monitoring is a new way to describe resistance (figure 5d), where an accumulative value of leaf area highlights the resistance of a genotype. Figure 5e and f show the development of *c*, where we observe similar properties for disease quantification as with *DS*_*l*_. The shape of instance should influence the relevance of *c* for disease quantification. In our past contribution [11], we calculate *c* from a circle shape instance and do not find relevance for disease quantification and variety differentiation for resistance. However, using leaf form instances from our instance segmentation model, we are able to contrast variety quantitative resistance.

**Figure 5:**
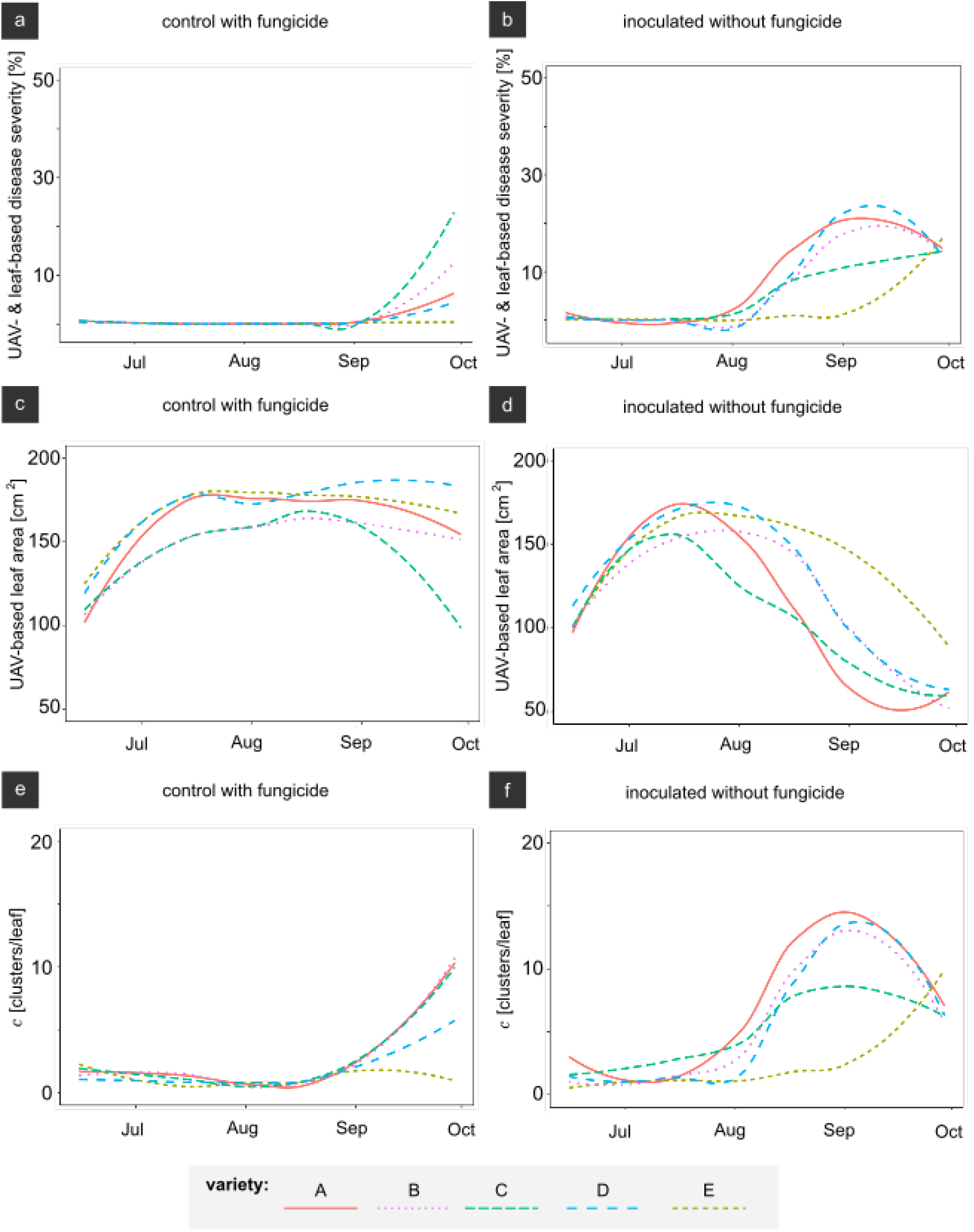
Development within fungicide strategies and five sugar beet varieties of leaf-based disease quantification parameters: disease severity (a and b), leaf area (c and d), and number of clusters per leaf (e and f). A total of 100 leaves were used for each treatment level. Curves were smoothed using locally estimated scatterplot method.

#### 2.2.7 Image-based disease incidence for decision making for plant protection

In Germany, thresholds of *DI* are the principal metric for decision making to control leaf diseases and to avoid economical losses in sugar beet cultivation [6]. On the field and for the case of CLS, leaves are ordered as “diseased” when at least one CLS spot is present on a sampled middle leaf. To implement an automatic scoring apporach, an accurate and robust detection of single spots is compulsory. However, current pixel-wise UAV-based image processing approaches are far from 100% level of precision and recall [9, 11]. This disadvantage makes the automatic definition of a diseased unit a challenge due to the risk of false positives if the criteria “*c* ≥ 1” is considered, because of the wrong classification of healthy regions as diseased increases. However, this criterium is easy to adapt to different disease quantification parameters in image-processing approaches (e.g. by using *c* or *ds*_*l*_) and diverse threshold values (Fig. 6a). We emulate possible time-series developments of *DI* with four different threshold values that defines a diseased unit (“diseased” *if* {*c* | *ds*_*l*_ ≥ 1, 5, 10, 25} number of clusters or %). In Fig 6b we observe that *DI* based on *c* parameter has a slightly advantage in comparison with *DI*s based on *ds*_*l*_ criteria, due to the early exponential phase in *DI* curve. Nevertheless, *DI* with the most sensible definition criteria of “diseased” (“diseased” if {*c* ≥ 1}) shows the disadvantage of the inaccurate pixel-wise approach, delivering high values of *DI* at the begin of July when the pathogen was just inoculated, with false positive detections. This inaccuracy can lead to a wrong decision for disease control. The alternative to this problem is to fix a higher threshold value to define a diseased unit avoiding possible emergence of false positives as observed in criteria {*c* ≥ 5, 10, 25}.

However, this decision shifts the exponential phase of *DI* to a later point in time, making this adaptation not suitable for practical use.

**Figure 6:**
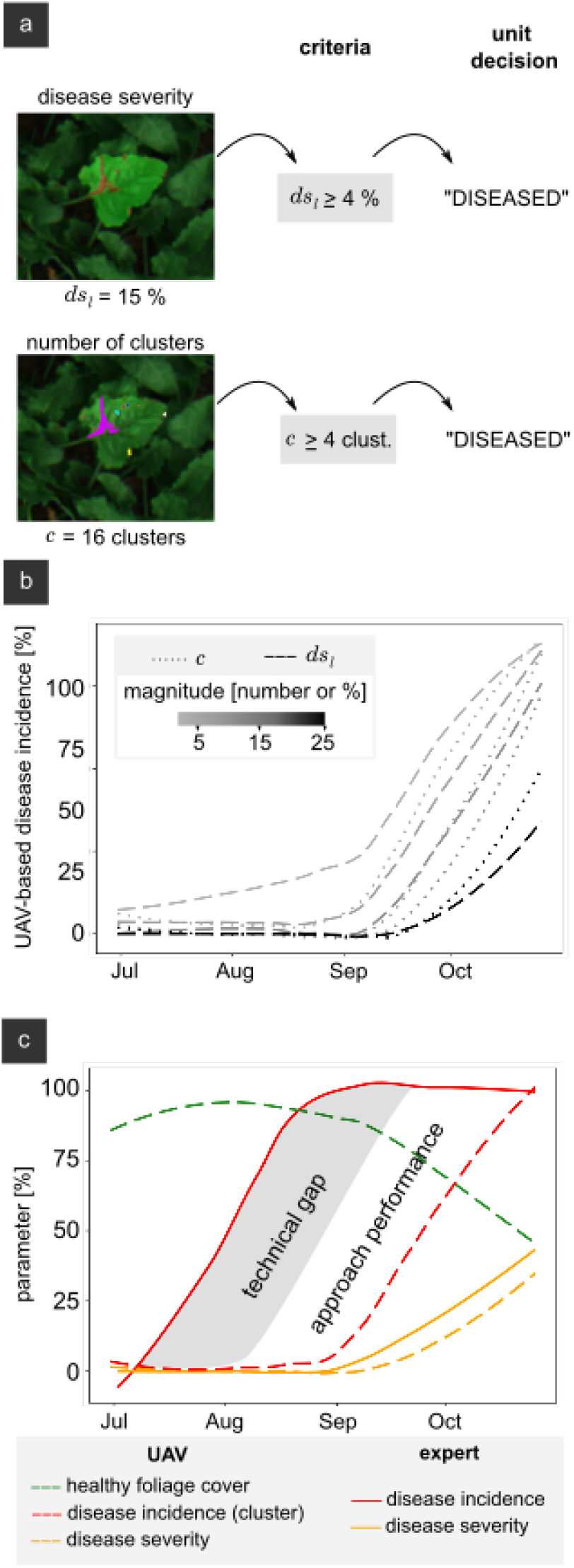
Decision making based on disease incidence (*DI*). a) Criteria for defining a diseased unit based by thresholding *c* or *ds*_*l*_ parameter. b) Development of *DI* curve based on diverse threshold magnitudes individually for *c* and *ds*_*l*_ parameters. c) Development of UAV-based and expert-based parameters. A total of 100 leaves were used for the treatment level in time-series (variety B and control with fungicide). Curves were smoothed using locally estimated scatterplot method.

In Fig 6c, we visualize the development of expert-based and UAV-based parameters. We observe that *DS* from expert-based and our leaf segmentation UAV-based data stay close together. This demonstrates that multispectral UAV-imaging for an automatic *DS* assessment under field conditions is feasible. However, in the case of *DI*, the expert ratings show an early exponential phase in the curve in July, while UAV-based scoring with a criteria “*c* ≥ 5” presents an exponential phase two months later than the experts. The performance of UAV-based *DI* parameter can be improved with a higher precision and recall for detecting diseased regions. However, we suspect there is a technical gap in the case of CLS for multispectral VISNIR UAV-imaging comparison to expert evaluation. We believe that this is due to restricted view of nadir perspective of UAV-based images in comparison to the possibility of the expert to sample middle and old sugar beet leaves individually, where first symptoms are observed [29].

### 2.3 Deep learning modeling and parameter extraction for phenotyping infested fields

In this work we give an overview of the complexity of modeling a leaf segmentation approach for UAV-images under field condition. For a practical use, an automatic leaf segmentation approach should provide robustness. This includes a good performance under diverse illumination conditions, for a high number of genotypes, and considering the epidemiological development of infested plants. This requires a high number of annotated images for modeling one plant-pathogen interaction.

Mask R-CNN models with a good AP performance deliver more accurate disease quantification parameters according to our results in comparison with models with a low AP. Leaf-based phenotyping parameters are relevant tools to describe epidemiological development of sugar beet genotypes under diverse disease pressure. Accumulated *A*_*l*_ over the complete monitoring season should differentiate genotypes for resistance against leaf diseases.

Instance form, whether circle or leaf, should influence the relevance of respective parameters to differentiate genotypes. To graph this statement, we evaluate the case for *c* parameter from our past contribution [11], where circlefrom *c* showed no significance for the genotype factor, while a leaf-form *c* presents a potential for resistance differentiation (Fig. 5c).

One of the principal limits for monitoring and scoring leaf diseases in field experiments is the necessity of a high number of trained personal. An automatic UAV-based scoring offers the chance to eliminate human error proper from individual scoring increasing the efficiency of collecting data.

The potential application of multispectral UAV-imaging and leaf-based parameters for disease management has to be analyzed in future studies. Although we mention the possible technical gap for UAV-based *DI*, we have to consider the potential of dividing a mapped field in mini plots for calculating *DI*, to make an individual and geo-referenced decision in large mapped fields.

## 3 Conclusion

We conclude that for leaf segmentation modeling of diseased sugar beet plants with Mask R-CNN, basic image augmentation techniques are more beneficial compared to a copy-paste approach. We demonstrate that a leaf-segmentation Mask R-CNN model can be integrated in a pipeline to extract leaf-based parameters of monitored fields. Although we evaluate only the case of CLS disease, the present pipeline for leaf-based parameter extraction can be transferred to agricultural practice and can support decision making in plant breeding for resistance and integrated pest management. Overall, we demonstrate that UAV-based monitoring of sugar beet fields followed by proper post-processing, can output reliable information that increases efficiency by replacing the very laborious work of visual scoring.

## Glossary

*A*_*D*_: surface area of diseased foliage within leaf instance.
*A*_*H*_: surface area of healthy foliage within leaf instance.
*A*_*L*_: image surface area within leaf instance.
*A*_*c*_: average cluster area within leaf instance.
*A*_*l*_: surface area within leaf instance.
*DI*: disease incidence.
*DS*: disease severity.
*DS*_*l*_: area based disease severity within leaf instance.
*L*: individual leaf instance.
ζ_*l*_: average slope or angle between surface and normal to horizontal within a leag instance.
ζ_*L*_: image slope or angle between surface and normal to horizontal within a leag instance.
*c*: number of clusters.
*ds*_*l*_: cover based disease severity within leaf instance.
AP: average precision.
CLS: Cercospora leaf spot.
CS: confidence score.
DSM: digital surface model.
GSD: ground sample distance.
IoU: intersection over union.
PLS-DA: partial least squares discriminant analysis.
ROI: region of interest.
SVMR: support vector machine radial.
UAV: unmanned aerial vehicle.

